# Cryo-EM structure and inhibitor design of human IAPP (amylin) fibrils

**DOI:** 10.1101/2020.05.25.114967

**Authors:** Qin Cao, David R. Boyer, Michael R. Sawaya, Peng Ge, David S. Eisenberg

## Abstract

Human islet amyloid polypeptide (hIAPP, or amylin) is a 37 amino acid hormone secreted by pancreatic islet β-cells. Aggregation of hIAPP into amyloid fibrils is found in more than 90% of Type-II Diabetes (T2D) patients and is considered to be associated with T2D pathology. Although different models have been proposed, the high resolution structure of hIAPP fibrils is unknown. Here we report the cryo-EM structure of recombinant full-length hIAPP fibrils. The fibril is composed of two symmetrically-related protofilaments with ordered residues 14-37 that meet at a 14-residue central hydrophobic core. Our hIAPP fibril structure (i) supports the previous hypothesis that residues 20-29, especially 23-29 are the primary amyloid core of hIAPP, (ii) suggests a molecular mechanism for the action of the hIAPP hereditary mutation S20G, (iii) explains why the 6 residue substitutions in rodent IAPP prevent aggregation, and (iv) suggests possible regions responsible for the observed hIAPP cross-seeding with β-amyloid. Furthermore, we performed structure-based inhibitor design to generate potential hIAPP aggregation inhibitors via a capping strategy. Four of the designed candidates delay hIAPP aggregation in vitro, providing a starting point for the development of T2D therapeutics and proof-of-concept that the capping strategy can be used on full-length cryo-EM fibril structures.

## Introduction

Formation of amyloid fibrils is associated with multiple diseases, including Aβ and tau with Alzheimer’s disease, α-synuclein with Parkinson’s disease (PD), and human islet amyloid polypeptide (hIAPP, or amylin) with Type-II Diabetes (T2D)^1^. hIAPP is a 37-residue hormone co-secreted with insulin by pancreatic islet β-cells to modulate glucose levels^2,3^. hIAPP has been identified as the major component of pancreatic amyloid deposits, which are present in more than 90% of T2D patients^4–6^. Several lines of evidence indicate that aggregation of hIAPP plays a key role in pathogenesis of T2D: i) the extent of islet amyloid positively correlates with pancreatic β-cell loss and insulin dependence^7–9^; ii) compared to humans, rodents do not normally experience T2D and do not form IAPP amyloid presumably because 6 residues differ between rodent and human IAPP^10,11^. However, mouse islet amyloid and T2D can be induced in mice by expression of hIAPP and a high fat diet^12,13^; iii) the hIAPP hereditary disease mutation S20G is associated with early onset T2D^14,15^, which correlates with its ability to accelerate hIAPP amyloid formation in vitro^16–18^; iv) hIAPP fibrils are toxic to pancreatic β-cells^19,20^.

The hIAPP segment containing residues 20-29 was initially proposed as the amyloid core because rodent hIAPP which does not form fibrils has 5 sequence differences in this segment^10^. Subsequent in vitro studies proposed additional residues within amyloid cores of hIAPP, including residues 12-17, 13-18, 15-20, and 30-37, whereas residues 1-13 were proposed not be involved in fibril formation since this segment cannot form fibrils by itself^21–24^.

Although two models of full-length hIAPP fibrils have been proposed based on experimental constraints^25,26^, the near-atomic resolution structure of hIAPP fibrils is still unknown. Here we determined the cryo-EM structure of full-length hIAPP fibrils at a resolution of 3.7 Å, and designed hIAPP aggregation inhibitors based on this structure.

## Results

### Construct design and fibril preparation

We generated full-length wild type human IAPP fibrils using a strategy similar to that reported previously^27^. We expressed and purified recombinant hIAPP with an N-terminal SUMO-tag. We intended to cleave off the SUMO-tag by growing the fibrils in the presence of ULP1 protease. However, we found that the SUMO-tag remained attached after fibril formation (see discussion). We pursued structure determination of these SUMO-tagged fibrils reasoning that hIAPP must compose the core of the fibrils and that SUMO is simply a bystander; our previous work showed that the SUMO-tag alone does not form aggregates^27,28^. Our observation that hIAPP can drive fibril formation even when tagged with a soluble, globular SUMO domain is consistent with previous studies that suggest hIAPP is one of the most amyloidogenic of amyloid-forming sequences^16^. After optimizing fibril growth and cryo-EM grid preparation, we were able to determine a 3.7 Å resolution cryo-EM structure of these SUMO-tagged hIAPP fibrils using helical reconstruction methods (Fig. 1, Table 1 and Supplementary Fig. 1).

**Figure 1.**
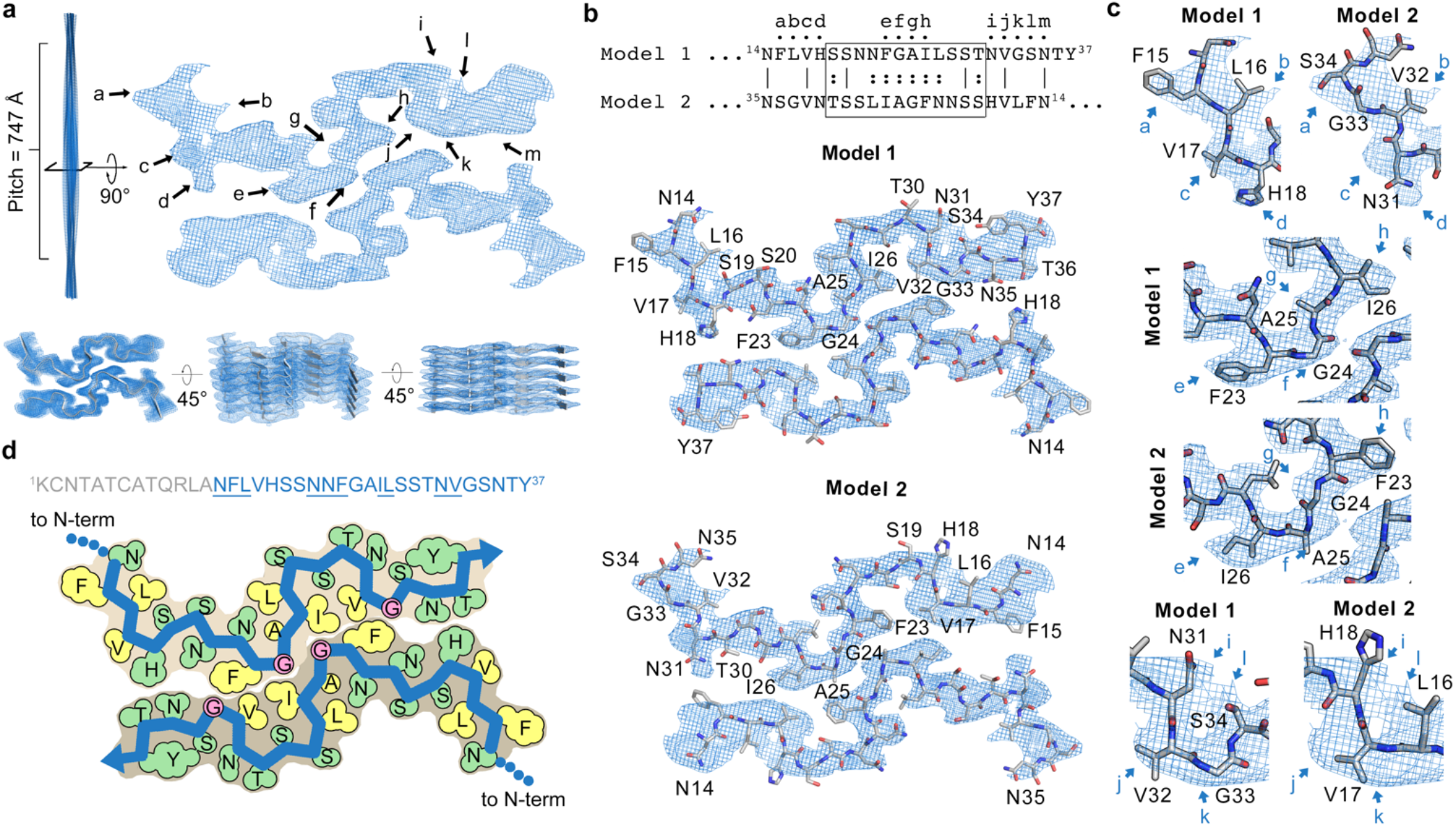
Cryo-EM map of hIAPP fibrils and identification of two models. **a,** (left) Side-view of fibril reconstruction, and (right) density of one layer of the fibril viewed down the fibril axis. (Bottom) different views of cryo-EM map with 5 layers shown. Top-view (left) demonstrates clear separation of β-strands while tilted views (middle, right) demonstrate clear separation of the layers of β-sheets along the fibril axis. b, Two models with opposite orientations were built and refined because of the quasi-symmetric nature of the hIAPP sequence, especially in the boxed region of the sequence alignment. The lines between two sequences represent identical residues and the colons represent similar residues. c, Detailed comparison of the density fit between two models. Positions used for identify the orientation of hIAPP models are labeled from a to m with arrows in panel a and c, and also above corresponding residues of Model 1 in panel b. d, Schematic view of Model 1, the protein backbone is colored blue, hydrophobic residues are colored yellow, polar residues are colored green, and glycine residues are colored pink. The amino acid sequence of hIAPP is shown above, with residues visible in the fibril structure colored blue and other residues colored grey. The residues adopting β-strand conformations are underlined.

**Table 1.** CryoEM data collection, refinement and validation statistics

|  |  |
| --- | --- |
|  | hIAPP<br>(EMD-21410,<br>PDB 6VW2) |
| **Data collection and processing** |  |
| Magnification | ×130,000 |
| Voltage (kV) | 300 |
| Electron exposure (e<sup>-</sup>/Å<sup>2</sup>) | 44 |
| Defocus range (μm) | 1.1-3.8 |
| Pixel size (Å) | 1.064 |
| Symmetry imposed | C<sub>1</sub> |
| Helical rise (Å) | 2.406 |
| Helical twist (°) | 179.42 |
| Initial particle images (no.) | 199,070 |
| Final particle images (no.) | 25,767 |
| Map resolution (Å) | 3.4 |
| FSC threshold | 0.143 |
| Map resolution range (Å) | 200-3.4 |
| **Refinement** |  |
| Initial model used (PDB code) | De novo |
| Model resolution (Å) | 3.7 |
| FSC threshold | 0.5 |
| Model resolution range (Å) | 200-3.7 |
| Map sharpening *B* factor (Å<sup>2</sup>) | 153.1 |
| Model composition |  |
| Nonhydrogen atoms | 1800 |
| Protein residues | 240 |
| Ligands | 0 |
| *B* factors (Å<sup>2</sup>) |  |
| Protein | 105.1 |
| Ligand | - |
| R.m.s. deviations |  |
| Bond lengths (Å) | 0.006 |
| Bond angles (°) | 1.138 |
| **Validation** |  |
| MolProbity score | 2.69 |
| Clashscore | 23.8 |
| Poor rotamers (%) | 0 |
| Ramachandran plot |  |
| Favored (%) | 72.7 |
| Allowed (%) | 27.3 |
| Disallowed (%) | 0 |

### Cryo-EM map of hIAPP fibrils

We found two fibril morphologies after performing 2D class-averaging of larger box sizes of 1024 pixels: one is a twisting fibril with an apparent two-fold symmetry axis that represents 78.4% of the identifiable particles and the other is a ribbon-like fibril that represents 21.8% of the identifiable particles (Supplementary Fig. 1a&b). As the structure of the ribbon is not readily determined by helical reconstruction because of its non-twisting nature, we focused on determining the structure of the twisting species.

Our map reveals the twister fibril is composed of two protofilaments related by a pseudo-21 axis (Fig. 1a). Pairs of hIAPP molecules adopt a flat, meandering topology, stacking as layers to assemble protofilaments. The layers are spaced 4.8 Å apart, hydrogen-bonded as parallel, in-register β-sheets – an arrangement similar to all previously determined cryo-EM amyloid fibril structures, such as Tau^29–31^, α-synuclein^32,33^, β2-microglobulin^34^ and TDP-43^27^.

### Identification of the model orientation of hIAPP fibrils

Establishing the orientation of the IAPP protein chain in our cryo-EM map posed a challenge because of the quasi-symmetrical amino acid sequence of the ordered core and the limited resolution of the map. Accordingly, we built and refined four atomic models and analyzed the compatibility of each with the cryo-EM map: Model 1 contains residues 14-37 and Model 2 contains residues 14-35 with opposite chain orientations (Fig. 1b, see Methods for detail). Both models fit the density equally well from Ser19 to Thr30 because the sequence of this segment is quasi-symmetric (see boxed region of sequence alignment in Fig. 1b). To determine which model fits the density better, we mainly rely on the fit of the sequence in the N- and C-terminal ends of the density as follows: i) On the left side of the top protofilament, there are four consecutive big bumps (labelled as position a-d in Fig. 1a&c), suggesting four consecutive big residues of the model, which agree with the sequence in Model 1 (Phe15-Leu16-Val17-His18). However, the corresponding sequence of Model 2 is Ser34-Gly33-Val32-Asn31, with two big residues and two small ones. Thus, Model 1 fits all four big bumps, while Model 2 does not (Fig. 1c, upper panel). In the best model we refined for Model 2, position c could not be fit and position d could be partially fit. ii) For position k, the density and space availability (close contact to the opposite protofilament) suggest a small residue, which agrees with Gly33 in Model 1. Model 1 is also supported by the kink in the mainchain density at this position, which is compatible with glycine. However, in Model 2 the corresponding residue is Leu16, which is too big for position k. To fit the density, we have to slide Leu16 into position l, which results in both Val17 and Leu16 of Model 2 only partially fitting into the mainchain density at positions j and l. In contrast, in Model 1, positions j and l are fully fitted by Val32 and Ser34 (Fig. 1c, bottom panel). iii) Positions i and m do not have big bumps, making His18 and Phe15 in Model 2 incompatible with the density, whereas in Model 1 these two positions are occupied by Asn31 and Asn35, which are smaller than His18 and Phe15 and consequently fit the density better. Further, His18 and Phe15 in Model 1 are well placed in positions d and a, respectively. iv) Gly24-Ala25 in Model 1 fits the density in position f and g better than in Model 2 (Fig. 1c, middle panel). v) The bump size of positions e and h also support Model 1 as the correct orientation: e is slightly bigger than h, so that Model 1 (Phe23 in e and Ile26 in h) offers a slightly better fit than Model 2 (Ile26 in e and Phe23 in h). We also analyzed the fit of two domain-swapped models, which consider the possibility of alternative connections of the two protofilaments. We found that Gly24 in the swapped models clearly does not fit the density, making the swapped models un-supported by our experimental map (Supplementary Fig. 2, see Methods for details).

In short, we believe that Model 1 is the better interpretation of the experimental map, meanwhile we note Model 2 fits the density fairly well and has acceptable peptide geometry and stability (Supplementary Table 1), so we cannot completely rule out the possibility that Model 2 is the correct structure of hIAPP. The ambiguity arises from the quasi-symmetric nature of the hIAPP sequence and the resolution limit of our cryo-EM map. We will use Model 1 as the preferred model of the hIAPP fibril for our following analysis.

### Cryo-EM structure of hIAPP fibrils

Our structure shows that residues 14-37 compose the fibril core. However, we see no density for the N-terminal residues 1-13 indicating they are dynamic. Similarly, we see no density for the SUMO-tag, which suggests that the SUMO-tag is also flexible and leads us to believe the SUMO-tag does not disturb the structure of the hIAPP fibril core (Supplementary Fig. 3, see discussion).

Hydrophobic forces strengthen the interface between molecules at the protofilament interface. Specifically, this interface is composed of side chains from Phe23, Ala25, Ile26, Leu27, and Val32, together with Gly24 and Gly33 (Fig. 1b, middle panel and Fig. 2a). Previous comparisons between pathogenic, irreversible fibrils structures and the fibril structure of FUS, which is considered to be involved in functional and reversible aggregation, suggested that the key feature determining the stability of the fibrils is the enrichment of hydrophobic residues buried in their fibril cores^27,35^. The 14-residue central hydrophobic core we observed suggests our hIAPP fibril structure is stable and likely irreversible – consistent with the pathogenic role of hIAPP fibrils in T2D. To attempt to quantify the stability of the hIAPP fibrils, we performed a solvation energy calculation similar to our previous studies^27,33^. The results are consistent with our qualitative analysis described above, and indicate that the central hydrophobic core is the most stable part of the fibril structure (Fig. 2b). The overall stability judged by energy per residue (−0.45 kcal/mol) is similar to previously reported irreversible fibril structures (i.e. −0.47 kcal/mol for TDP-43 SegA-sym^27^, −0.42 kcal/mol for Aβ ex vivo^36^, and −0.43 kcal/mol for tau PHF^29^), and much larger in value than FUS^35^ (−0.20 kcal/mol, see Supplementary Table 2). These results support the hypothesis that the hIAPP fibrils reported here are essentially irreversible.

**Figure 2.**
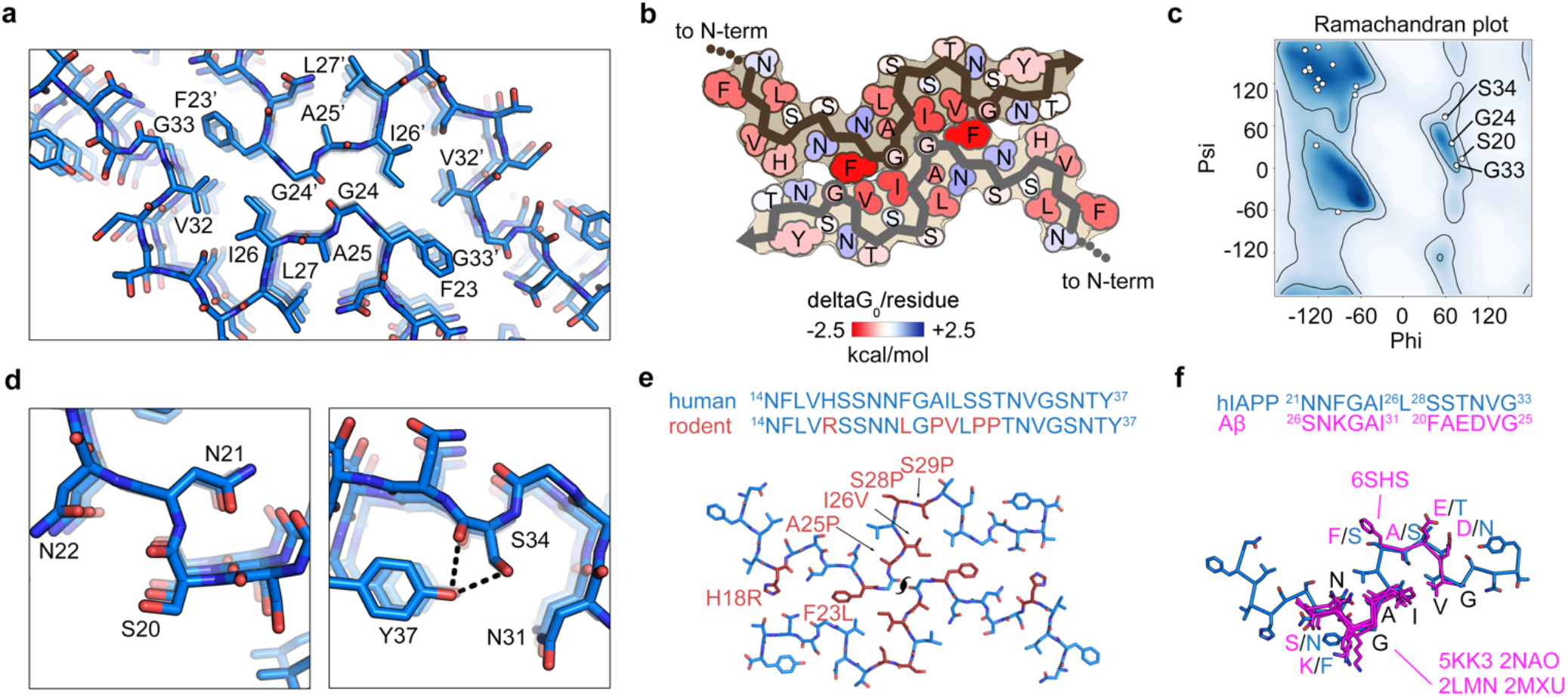
Structure analysis of hIAPP fibrils. **a,** The central hydrophobic core that stabilizes the hIAPP fibril structure. **b,** Solvation energy map of hIAPP fibril structure. Residues are colored from unfavorable (blue, 2.5 kcal/mol) to favorable stabilization energy (red, −2.5 kcal/mol). **c,** Ramachandran plot of hIAPP fibril structure, labeling residues having positive Phi angles. **d,** Hydrogen bonding networks around Ser20 (left) and Ser34 (right). Hydrogen bonds with distance between 2.3-3.2 Å are shown as black dashed lines. **e,** Comparison of human and rodent IAPP sequence in the region that is visible in hIAPP fibril core. The residues that differ between rodent and human are colored in red. Notice that the human residues all contribute to stabilizing the hIAPP fibril structure, so that the rodent substitutions are expected to disrupt fibril formation. **f,** Alignment of previously reported Aβ fibril structures (magenta, see Methods) with hIAPP fibril structure (blue) suggests two pairs of possible cross-seeding regions. For detailed alignment parameters see Supplementary Fig. 5 and Supplementary Table 3.

### Possible mechanism of the S20G hereditary mutation in Type 2 diabetes patients

Our hIAPP fibril structure suggests the molecular mechanism by which the hIAPP hereditary mutation, S20G, accelerates fibril growth in vitro^16–18^, and thus perhaps in vivo. The finding of S20G in a small fraction of younger Japanese type 2 diabetes patients may suggest that the S20G variant forms more stable fibrils than the wild type sequence^37^. In fact our structure suggests that glycine in position 20 confers greater fibril stability than serine. Ser20 in our hIAPP fibril structure adopts a positive ϕ-backbone dihedral angle, placing it in the left-handled alpha-helix region of the Ramachandran diagram (Fig. 2c). Sidechains other than the hydrogen atom of glycine tend to suffer various degrees of steric hindrance in this region. Hence the replacement of serine by glycine would relieve the steric hindrance and favor fibril formation. Ser34 also adopts a positive ϕ-backbone dihedral angle in our model (Fig. 2c) but this strained conformation is stabilized by a hydrogen bond between the hydroxyl group of Ser 34 and Tyr37. Ser20 lacks an analogous stabilizing interaction (Fig. 2d). These observations may explain why S20G is a disease-related mutation of hIAPP but S34G is not.

### Rodent substitutions disrupt hIAPP fibril structure

Our hIAPP fibril structure also explains why rodent IAPP (rIAPP) does not fibrillize as readily as hIAPP^38^. Our structure shows that residues 23-29 comprise the fibril core of hIAPP, consistent with previous predictions that these residues are essential for aggregation. Five out of the six amino acid differences between human and rodent IAPP are located here. Notably, four residues in this segment contribute the majority of the hydrophobic forces that stabilize fibril assembly (Phe23, Gly24, Ala25 and Ile26, Fig. 2e). The only residue difference between hIAPP and rIAPP that is located outside this core is His18; it is oriented to form a water-mediated hydrogen bond with Thr36 from the opposite protofilament (Fig. 1b, middle panel). Taken together, all 6 residues that differ between hIAPP and rIAPP contribute fundamentally to the stability of the hIAPP fibril core; hence, we expect that substitutions of these residues would seriously destabilize the fibril structure (Fig. 2e).

To quantify how destabilizing the rat sequence would be to fibril formation, we modeled rodent substitutions H18R, A25P, I26V, S28P, and S29P computationally on our hIAPP fibril structure. We found steric clashes in several areas that could not be relieved by alternative side chain rotamers. The Rosetta energy remained strongly unfavorable (+3549 REU, Rosetta energy units, Supplementary Fig. 4a) even after side chain repacking. The primary clashes in the rodent model occurred at S28P and S29P, accompanied by the elimination of two backbone hydrogen bonds (Supplementary Fig. 4b). Rosetta energy minimization could accommodate these substitutions only by breaking apart the fibril fold, imposing a r.m.s. deviation from the starting hIAPP model of 5.4 Å (Supplementary Fig. 4a). In contrast, the Rosetta energy for the corresponding human model is favorable, - 41 REU per chain (Supplementary Fig. 4a).These results indicate that the rIAPP sequence is fundamentally incompatible with our hIAPP fibril structure and consistent with the observation that mice and rats do not form islet amyloid fibrils.

### Comparison between hIAPP and Aβ fibril structures

Recent studies suggest a correlation between T2D and AD via potential cross-seeding between aggregates of hIAPP and β-amyloid (Aβ), a 39-42 residue peptide that plays a central role in AD pathogenesis. hIAPP has been found to co-deposit with Aβ in senile plaques in AD patient-derived samples. Increased risk for AD in T2D patients (and vice versa) has been reported by clinical studies^40–42^. Cross-seeding effects between hIAPP and Aβ have been found in vitro and in transgenic mice^39,43–45^. The cross-seeding between hIAPP and Aβ might be due to the structural similarity between aggregates of these two proteins whereby aggregates of one protein may be able to template the formation of aggregates of the other. Our previous study proposed a model where the cores responsible for cross-seeding are hIAPP 19-29 and Aβ 24-34, which share 50% sequence similarity^45^. We superimposed our hIAPP fibril structure with previously reported Aβ fibril structures, either by aligning whole structures without restrictions, or by restricting the alignment to Aβ 24-34 and hIAPP 19-29, which follows our previous prediction of cross-seeding core. By whole structure superimposing, we found one of the Aβ structures, Aβ ex vivo fibril structure (PDB ID 6SHS), contains a 6-residue region, Aβ 20-25, that shares a similar conformation with hIAPP 28-33 with an overall r.m.s.d. of 1.8 Å (Fig. 2f, Supplementary Fig. 5 and Supplementary Table 3). By restricted superimposing, we find another four Aβ structures show high structure similarity with our hIAPP structure at Aβ 26-31 and hIAPP 21-26, and three additional Aβ structures show partial similarity at the same area, with r.m.s.d. all below or equal to 2 Å (Fig. 2f, Supplementary Fig. 5 and Supplementary Table 3). Taken together, our hIAPP fibril structure supports the potential for cross-seeding between hIAPP and Aβ and suggests two potential cores responsible for this effect, which may further provide information for Hot-Segment-Based inhibitor design^46^.

### Structure-based inhibitor design of hIAPP fibrils

Our previous studies demonstrated that by designing peptides that bind to fibril cores and sterically prevent further addition of monomer to the fibril ends, we are able to develop capping inhibitors that block amyloidogenesis^47–51^. This strategy has been demonstrated successfully using crystal structures of peptide segments as templates for inhibitor design, but to our knowledge no inhibitors have been designed solely on full length fibril structures. Here we designed 6 inhibitors based on our hIAPP fibril structure targeting three regions, ^21^NNFGAILSS^29^ (N9S), ^25^AILSSTNVG^33^ (A9G) and ^21^NNFG^24^ (N4G). N9S and A9G are both 9 residue segments from a single protofilament, whereas 2 copies of N4Gs from both protofilaments are merged into one peptide (N4Gm) to cap two protofilaments simultaneously (Supplementary Fig. 6). Because N4Gs from two protofilaments in the hIAPP fibril structure are facing each other with opposite N-terminal to C-terminal orientations, in order to generate a peptide with the correct orientation, we reversed the sequence of one N4G segment before linking it with another (Supplementary Fig. 6). We also reversed the chirality of the N4G segment whose sequence was reversed to mimic the conformation of the target N4G in the fibril structure so that the merged peptide can bind both protofilaments simultaneously (Supplementary Fig. 6 and Supplementary Table 4, see Method for detail). We then added main chain N-methylation to the designed peptide to block hydrogen bond formation with additional hIAPP molecules, resulting in 6 inhibitor candidates (Fig. 3a, Supplementary Fig. 7a, and Supplementary Table 4). We tested the inhibition of hIAPP fibril growth using these candidates with ThT aggregation assays and found four candidates delay aggregation of hIAPP (Fig. 3b and Supplementary Fig. 7b). Negative stain EM images suggest the successful inhibitors reduced the amount of fibrils that form, consistent with our ThT measurements, whereas the existence of fibrils indicates the fibril formation is not fully eliminated nor re-directed into other aggregation pathways such as the formation of amorphous aggregates (Fig. 3c and Supplementary Fig. 7c). We note our inhibitors inhibit hIAPP fibril formation with low efficiency, since they delay but not eliminate hIAPP fibril formation, and mostly work only at a high molecular ratio (IAPP:inhibitor = 1:30). Nonetheless, we believe these inhibitors serve as a starting point for future inhibitor design and a proof-of-concept that the capping strategy works on full-length fibril structures. Further studies are needed to increase the efficiency of these inhibitors and develop potential leads for the therapeutics of T2D.

**Figure 3.**
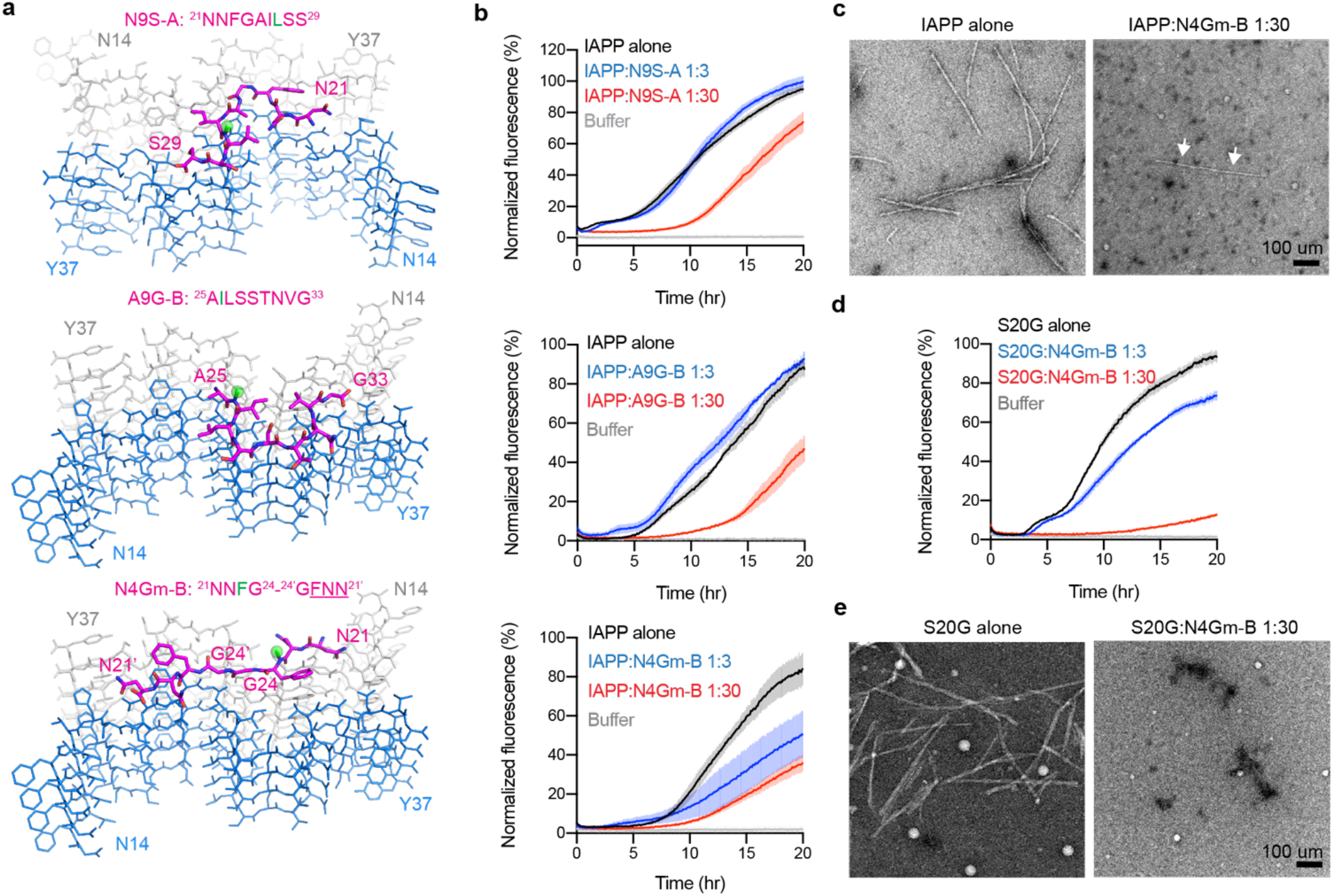
Structure-based inhibitor design of hIAPP fibrils. **a,** Proposed models of designed inhibitors (magenta) capping hIAPP fibrils. The methyl group of N-methylated inhibitors is shown as a green sphere. Notice N9S-A and A9G-B bind to a single protofilament, whereas N4Gm-B binds to both protofilaments. The last three residues of N4Gm-B are underlined and are D-amino acids. **b,** ThT assays demonstrate inhibitors delay hIAPP fibril formation. **c,** Negative stain EM images of hIAPP with or without N4Gm-B after 20 hours of incubation. Notice that hIAPP fibril formation is not fully eliminated by N4Gm-B (indicated by white arrows). **d-e,** ThT (**d**) and EM (**e**) demonstrate N4Gm-B inhibits fibril formation of hIAPP S20G. ThT data are shown as mean ± s.d., n=3 independent experiments.

Our inhibitor designs, especially N4Gm-B, can also be used as a probe to detect the existence of our hIAPP structure during hIAPP aggregation. Since N4Gm is designed based on the model that two N4Gs are close to each other with a face-to-face orientation, this inhibitor should be specific to the structure we determined here, not to other potential polymorphs. In the inhibition assays shown in Fig. 3, we used chemically synthesized hIAPP without SUMO-tag and with C-terminal amidation (see Discussion). The observation that N4Gm-B inhibits synthetic hIAPP fibril formation supports our hypothesis that the SUMO-tag and C-terminal amidation does not disturb the fibril structure we reported here given that the inhibitor N4Gm-B would not work on a different fibril structure. As we proposed previously, S20G favors hIAPP fibril formation by the adoption of a positive ϕ-backbone dihedral angle that is favorable for glycine which further suggests that hIAPP S20G should form the same structure as wild type hIAPP. We tested this hypothesis using our N4Gm-B probe, and indeed we found that N4Gm-B also inhibits hIAPP S20G aggregation via ThT assays and EM images (Fig. 3d&e).

## Discussion

The hIAPP protein we used for fibril structure determination has two major differences from hIAPP found in patient islet deposits: i) lack of C-terminal amidation and ii) presence of the SUMO-tag. We believe that neither of these differences invalidates the structure we determined here. A previous study demonstrated that regardless of C-terminal amidation state, hIAPP forms aggregates in vitro with similar kinetics, cytotoxicity, and fibril morphology^52^. Regarding the SUMO-tag, our cryo-EM reconstruction reveals a fuzzy coat of density outside the hIAPP fibril core (Supplementary Fig. 3a). Since the density is smeared in a semi-spherical pattern and centered near Asn14, we believe the density originates from the flexible N-terminal region of our hIAPP construct, which includes hIAPP 1-13 and the SUMO-tag. Although the SUMO-tag was designed to be cleaved during fibril formation, it remained attached likely due to the limited accessibility of protease to the linker between SUMO and hIAPP (Supplementary Fig. 3b). The dimensions of the density (radius ≈ 55 Å) fit the SUMO fold (35 Å x 26 Å x 18 Å) (Supplementary Fig. 3d, PDB ID 1L2N^53^) in addition to the 13-residue flexible N-terminus of hIAPP (Supplementary Fig. 3c, see next paragraph for detail). The fit suggests that hIAPP 1-13 and SUMO-tag freely wiggle around the fibril core with Asn14 acting as an anchor (Supplementary Fig. 3e). We expect contact between the SUMO-tag and the hIAPP fibril core to be weak and nonspecific since there are 13 flexible residues separating SUMO from the hIAPP N-terminus. Hence, the SUMO-tag should not disturb the core structure of hIAPP fibrils. This hypothesis is supported by the lack of ordered density around the hIAPP fibrils. Taken together, we believe that the lack of C-terminal amidation and the existence of a SUMO-tag should not influence the fibril structure we report here, which is also supported by the observation that the inhibitors designed based on our fibril structure inhibit fibril formation of synthetic IAPP peptides lacking a SUMO-tag and having C-terminal amidation.

In our full-length hIAPP fibril structure, we observed residues 1-13 are flexible and lack ordered density. This flexibility of the N-terminus is consistent with previous studies that mapped the amyloid cores of hIAPP to segments spanning the region 12-37, but not 1-13^10,21–23^. We also found smeared density that may correspond to the flexible 1-13 segment of hIAPP and the SUMO-tag, and within this region we observed additional density with a shape of a potential β-strand using a lower sigma cutoff (Supplementary Fig. 3c). We propose a plausible model of hIAPP 5-11 occupying these densities (see Methods) indicating a hypothetical preferred conformation of the flexible N-terminus of hIAPP. Whereas we believe this conformation of the N-terminus is not important for hIAPP fibril formation based on the previous knowledge of the segments necessary for fibril formation, and also based on the observation that these additional weak densities are only visible under a low noise cutoff and are located in a position that does not suggest any close contacts with the fibril core except with Phe15 in the current model (Supplementary Fig. 3c).

Two models of hIAPP fibrils have been previously proposed based either on NMR or EPR constraints^25,26^, and our structure differs from both of them. In addition, during the final preparation of our paper, Röder et al. deposited a preprint in BioRxiv titled “Amyloid fibril structure of islet amyloid polypeptide by cryo-electron microscopy reveals similarities with amyloid beta.” They report the structure of a polymorph of hIAPP called PM1, having two S-shaped protofilaments also with zipper-like bonding at the sequence NFGAIL, but with a different fold than ours. We note that pathogenic amyloid fibrils are usually polymorphic and the other models may therefore represent different polymorphs of hIAPP than the structure we determined here. In this case, since we observe that the S20G hereditary mutation may benefit the formation of our structure, we hypothesize that our structure may represent a polymorph preferred by the S20G mutant sequence.

In summary, the cryo-EM structure of hIAPP fibrils we reported here visualizes the amyloid core of hIAPP, offers a potential mechanism of the S20G hereditary disease mutation and aggregation-disrupting rodent mutations, suggests possible regions responsible for Aβ-hIAPP cross-seeding, and serves as a template for structure-based inhibitor design.

## Methods

Methods and materials used in this study are available in supplementary information.

## Acknowledgements

We thank H. Zhou for use of Electron Imaging Center for Nanomachines (EICN) resources. We acknowledge the use of instruments at the EICN supported by NIH (1S10RR23057 and IS10OD018111), NSF (DBI-1338135) and CNSI at UCLA. The authors acknowledge NIH AG 054022, NIH AG061847, and DOE DE-FC02-02ER63421 for support. D.R.B. was supported by the National Science Foundation Graduate Research Fellowship Program.

## Author contributions

Q.C. designed experiments, purified constructs, prepared cryo-EM samples, performed cryo-EM data collection and processing, designed inhibitors, performed biochemical experiments and performed data analysis. D.R.B. and P.G. assisted in cryo-EM data collection and processing. Q.C. and M.R.S. built the inhibitor binding model. M.R.S. performed solvation energy calculation. All authors analyzed the results and wrote the manuscript. D.S.E. supervised and guided the project.

## Materials & Correspondence

For requests of materials reported in this study, please contact David S. Eisenberg.

## Data availability

Structural data have been deposited into the Worldwide Protein Data Bank (wwPDB) and the Electron Microscopy Data Bank (EMDB) with accession codes: PDB 6VW2, EMD-21410. All other data, including the custom software used for solvation energy calculation, are available from the authors upon reasonable request.

## METHODS

### Construct design

SUMO-hIAPP constructs were designed with the same strategy as reported previously^1,2^, which uses a (His)_6_-tag for Ni-column purification, SUMO protein as a solubility tag and full-length hIAPP. hIAPP cDNA was inserted into a pET28a vector containing an amino terminus conjugation of SUMO protein. We initially designed one glycine residue as a linker between SUMO and hIAPP, and we found the SUMO tag cannot be cleaved from this construct. We then extended the linker to three glycine residues to generate a SUMO-tag removable construct. The sequences of both SUMO tag unremovable version (one-glycine linker, 1xG) and removable version (three-glycine linker, 3xG) are shown as follows:

SUMO-hIAPP (1xG)
MGSSHHHHHHGSGLVPRGSASMSDSEVNQEAKPEVKPEVKPETHINLKVSDGSSEIFFKIKKTTPLRRLM EAFAKRQGKEMDSLRFLYDGIRIQADQTPEDLDMEDNDIIEAHREQI**G**KCNTATCATQRLANFLVHSSNN FGAILSSTNVGSNTY

SUMO-hIAPP (3xG)
MGSSHHHHHHGSGLVPRGSASMSDSEVNQEAKPEVKPEVKPETHINLKVSDGSSEIFFKIKKTTPLRRLM EAFAKRQGKEMDSLRFLYDGIRIQADQTPEDLDMEDNDIIEAHREQI**GGG**KCNTATCATQRLANFLVHS SNNFGAILSSTNVGSNTY

### Protein purification and SUMO-tag cleavage assays

Protein purification follows the same protocol reported previously^1,2^. Both 1xG and 3xG SUMO-hIAPP were expressed in the Escherichia coli BL21 (DE3) strain, which grows in LB media with 50 μg/ml kanamycin. Protein expression was induced by adding 1 mM isopropyl β-D-1-thiogalactopyranoside (IPTG) to the cell culture when an OD600 of 0.6-0.8 was reached. The bacterial cells were further cultured at 25 °C for 3 hours and then were harvested and resuspended in 20 mM Tris-HCl, pH 8.0, 500 mM NaCl, 20 mM imidazole and 10% (v/v) glycerol, supplemented with 1% (v/v) halt protease inhibitor single-use cocktail (Thermo Scientific), and sonicated (3 s on/3 s of cycle, 10 min) and centrifuged (24,000g for 20 min) to obtain the cell lysate. We added our homemade NucA nuclease (5000 U per liter of cell culture) to the cell lysate, filtered the mixed solution and then loaded it onto a HisTrap HP column (GE healthcare). The column was pre-equilibrated with 20 mM Tris-HCl, pH 8.0, 500 mM NaCl and 20 mM imidazole before loading the sample, and after the sample was loaded, the column was washed with 20 mM Tris-HCl, pH 8.0, 500 mM NaCl and 200 mM imidazole and eluted with 20 mM Tris-HCl, pH 8.0, 500 mM NaCl and 500 mM imidazole. Purified protein was concentrated using Amicon Ultra-15 centrifugal filters (Millipore) and stored at −80 °C for future use.

To remove the SUMO-tag, ULP1 protease cleavage assays were performed as reported previously^61,2^. Both 1xG and 3xG SUMO-hIAPP protein were mixed with 100:1 (weight basis) with homemade ULP1 protease and samples were analyzed via SDS-PAGE (Supplementary Fig. 3b). At 0 h, samples mixed with ULP1 showed bands of intact SUMO-hIAPP; at 1h of cleavage, only 3xG SUMO-hIAPP showed band of free SUMO and hIAPP, whereas 1xG still showed intact SUMO-hIAPP, indicating 3xG is SUMO removable but 1xG is not; after more than one month (> 1 m) of incubation at fibril growth condition, SUMO-hIAPP form fibrils and still showed intact SUMO-hIAPP band but not free SUMO and hIAPP bands, indicating 1xG SUMO-hIAPP fibrils contain SUMO tags.

### Synthetic peptide preparation

Full-length hIAPP wild type and S20G were synthesized by InnoPep with amination at the carboxyl terminal and an intramolecular disulfide bridge between Cys2 and Cys7 with a purity higher than 95%.The inhibitors were synthesized by GenScript at a purity of 95% or higher. The sequences of all inhibitors designed are summarized in Supplementary Table 4, and the sequences of wild type and S20G hIAPP peptides are as follows:

hIAPP wild type:
KCNTATCATQRLANFLVHSSNNFGAILSSTNVGSNTY-NH_2_

hIAPP S20G:
KCNTATCATQRLANFLVHSGNNFGAILSSTNVGSNTY-NH_2_

The synthetic hIAPP peptide was first dissolved in 100% HFIP at a concentration of 1 mM, sonicated at 4 °C for 1 min, and incubated at room temperature for 5 hours. The HFIP was then removed with a CentriVap Concentrator (Labconco) and treated peptides were stored at −20 °C. Before use, the peptides were freshly dissolved at 1 mM or 5 mM in 100% DMSO, and further diluted 100-fold in PBS and filtered using 0.1 μm Ultrafree-MC-VV centrifugal filters (Millipore) to form 10 μM and 50 μM hIAPP solutions. The peptide inhibitors were dissolved in 100% DMSO at a concentration of 30 mM and stored at −20 °C. Before use, the DMSO inhibitor stocks were diluted to 10 mM or 3 mM with 100% DMSO, and then diluted 100-fold into hIAPP PBS solution to form 300 μm, 100 μm and 30 μm inhibitor mixtures.

### Fibril preparation and optimization

Recombinant 1xG SUMO-hIAPP protein was diluted to 50 uM with 20 mM Tris-HCl, pH 8.0, 500 mM NaCl, and mixed with 100:1 (weight basis) homemade ULP1 protease. Fibril formation was inspected using negative-stain transmission election microscopy after 3 days of shaking at 37 °C. In order to obtain the fibrils suitable for cryo-EM data collection, the fibril growth condition was optimized by modifying protein concentration, incubation time and temperature, buffer (pH, salt concentration, additives) and seeding with preformed fibrils. The final fibril growth condition used for cryo-EM structure determination was 50 uM protein mixed with 100:1 ULP1 in 20 mM Tris-HCl, pH 8.0, 500 mM NaCl supplemented with 200 mM Imidazole and 100% (molar basis, monomer equivalent) pre-formed fibril seeds and incubated at 37 °C without shaking. The pre-formed fibril seeds were formed with the same optimized condition but 2% (molecular basis, monomer equivalent) fibrils that formed with the initial fibril growth condition. The 2% seeded fibrils were concentrated via centrifugation at 4,000 g for 3 minutes, and sonicated at 4 °C for 30 minutes to ensure the complete fragmentation of the fibrils, and the sonicated seeds were mixed with fresh SUMO-hIAPP monomers at a 100% molar basis for final fibril growth. The 100% seeded fibrils were used for cryo-EM sample preparation. We note that although ULP1 was added in the initial and optimized fibril growth conditions, the SUMO-tag was not cleaved from the constructs because of the 1xG linker (Supplementary Fig. 3b), so that the fibrils we used for cryo-EM structure determination contains SUMO-tag.

### Negative-stain transmission election microscopy

Negative-stain transmission EM samples were prepared by applying 5 μl of solution to 400 mesh carbon-coated formvar support films mounted on cooper grids (Ted Pella, Inc.). The grids were glow-discharged for 30 seconds before applying the samples. The samples were incubated on the grid for 2 minutes and then blotted off with a filter paper. The grids were stained with 3 μl of 2% uranyl acetate for 1 minute, and washed with an additional 3 μl of 2% uranyl acetate and allowed to dry for 1 minute. The grids were imaged using a T12 (FEI) election microscope.

### Cryo-EM data collection and processing

Cryo-EM data was collected with the same strategy reported previously^2^. A Quantifoil 1.2/1.3 electon microscope grid was glow-discharged for 4 minutes and applied with 2.8 μl of fibril solution. The grid was plunge frozen into liquid ethane with a Vitrobot Mark IV (FEI). Data were collected on a Titan Krios (FEI) microscope with a Gatan Quantum LS/K2 Summit direct electron detection camera. The microscope was operated with 300 kV acceleration voltage and slit width of 20 eV. Super-resolution movies were collected with a nominal physical pixel size of 1.07 Å/pixel (0.535 Å/pixel in super-resolution movie frames) with a dose per frame of ~1.1 e Å^2^ and dose per image of ~44 e^-^/Å^2^. Forty frames were taken for each movie with a frame rate of 5 Hz. Automated data collection was driven by the Leginon automation software package^3^. After data collection, anisotropic magnification distortion estimation, CTF estimation and beam-induced motion correction were performed with mag-distortion-estimate^4^, CTFFIND 4.1.8^5^ and Unblur^6^, respectively. The physical pixel size was corrected to 1.064 Å/pixel after anisotropic magnification correction with Unblur^6^.

Particle picking was performed manually using EMAN2 *e2helixboxer.py*^1^ with all fibrils picked as one group. Particles were extracted in RELION^8,9^ using the 90% overlap scheme into 1,024 and 686 pixel boxes, respectively. 2D Classification, helical reconstruction, and 3D refinement were also performed in RELION. We performed 2D classification with 1,024 and 686 pixel particles to identify the fibril morphologies, and found among all the particles that belong to 2D classes with recognizable features, ~78.4% particles belong to 2D classes that show twister morphology and ~21.6% particles belong to classes that show ribbon morphology (Supplementary Fig. 1b). We did not pursue 3D reconstruction with the ribbon morphology because it lacks twisting. We selected the 686 pixel particles from the twister morphology and performed 3D classification with the helical parameters estimated from the measured crossover distance and an elongated Gaussian blob as an initial model. After that, we ran an additional 3D classification with K=3 and used the preliminary reconstructions as a reference to select for particles contributing to homogenous classes with stable helicity and separation of β-strands in the x-y plane. To achieve this, we manually control the tau_fudge factor and healpix to reach a resolution of ~7 Å for the best class and the particles from this class were selected for the next step of refinement. To get the near-atomic resolution reconstruction, we applied a “re-extraction” strategy. We reran particle extraction using 90% overlap scheme with 288 pixel boxes, extracting particles only from the fibril tubes that had particles that were included in the best class of the previous 3D classification. The re-extracted particles were used for two rounds of a K=3 class 3D job to select the particles that contributed to the highest-resolution class. The selected particles were re-extracted again with CTF phase flipping and used for high-resolution gold-standard refinement as described previously^9^. For overall resolution estimation, we use the 0.143 FSC resolution cutoff which reports an estimated resolution of 3.4 Å. The reconstruction was compared against the reference-free 2D class averages to confirm its validity. In addition, de novo model building also confirmed the validity of the 3D reconstruction.

### Atomic model building

The refined map was sharpened via phenix.auto_sharpen^10^ at the resolution cutoff of 3.4 Å indicated by half-map FSC. The atomic model of the hIAPP fibril was manually built into the refined map using COOT^11^. We first inspected the density at the center of the dimer interface and found the pattern of the model should be XXxxXX based on the sizes of side chains where X stands for a large side chain residue and x stands for a small side chain residue. We inspected the sequence of hIAPP and identified ^22^NFGAIL^27^ is the best fit of this pattern. Our earlier x-ray diffraction study identified ^21^NNFGAIL^27^ as a stabilizing segment of hIAPP fibrils^12^. We therefore built the ^22^NFGAIL^27^ model into the density with both orientations. Further model building generated two possible models (Model 1 and 2, Fig. 1b) corresponding to the two orientations of the protein chain. Examination of weak density to identify the location of the flexible N-terminal residues did not help to choose between these models. The reason is that in the final map, the extra density seams to connect to the N-terminus of Model 2 (Supplementary Fig. 8, left panel); whereas in our earlier reconstruction at lower resolution which better shows the trace of mainchain, the extra density seams to connect to the N-terminus of Model 1 (Supplementary Fig. 8, right panel). So we refined both models and chose the preferred model based on their density fit (see results).

We also found that in both Model 1 and 2, Gly24 and Ala25 from both protofilament are close to each other and are in the center of the dimer interface (Supplementary Fig. 2). This observation suggests a possibility of domain swapping, which means the residues between the N-terminus to Gly24 from one protofilament can connect to residues between Ala25 to the C-terminus from the other protofilament. To test this possible domain swap, we built swapped version of both Model 1 and Model 2. We found although the rest of the residues of swapped models fit the density equally well with un-swapped models, the Gly24 residues in the swapped models are clearly out of the density, even with lower sigma (Supplementary Figure 2). So that we believe our density does not support the domain swap of our hIAPP models.

The initial models were extended to five layers (ten chains) based on the helical symmetry of the reconstruction using our in-house script^2^. The five-layer model was then refined by phenix.real_space_refine^13^. After several rounds of refinement, we adjusted the orientation of the main chain oxygen and nitrogen to facilitate main chain hydrogen bonding within the β-sheet, and we applied these main chain hydrogen bond restraints for further real-space refinement. As the last step, the rotamer of each serine, glutamine and asparagine residue was manually inspected to ensure energetically favorable hydrogen binding. The final model was validated using MolProbity^14^.

The final model fits well with the map for all side chains, and the map-to-model resolution was estimated to be 3.7 Å with 0.5 FSC cutoff. The observation that Asn14 to the C-terminus of hIAPP was forming the fibril core also agrees with previous studies that the hIAPP amyloid core is formed by the C-terminal region and residues 1-13 alone cannot form fibrils^15^. On both sides of the fibril core, we inspected additional densities apparent with a lower sigma cutoff of the map (Supplementary Fig. 2c). We modeled a seven-residue idealized β-strand into the densities without further modification because we cannot identify its registration, orientation and side chain conformation based on the quality of the density. We hypothesized it to be the N-terminal part of hIAPP based on its position and feasibility of the Cys2-Cys7 intramolecular di-sulfur bond. In this model, residues 5-11 occupy these extra densities that are 2 residues away from the first visible residue Asn14 and with Cys7 pointing away from the fibril core to form a di-sulfide bond with Cys2 (Supplementary Fig. 2c). We note that this model of 5-11 is only one of numerous plausible models, therefore we did not include it in our final structure. The low quality of these densities indicates the conformation of the N-terminus is flexible.

### Energetic calculation

The free energy is an adaptation of the solvation free energy described previously^16,17^, in which the energy is calculated as the sum of products of the area buried of each atom and its corresponding atomic solvation parameter (ASP). ASPs were taken from previous work^24^. Area buried is calculated as the difference in solvent accessible surface area (SASA) of the reference state (i.e. unfolded state) and the SASA of the folded state. The SASA of residue i of the unfolded state was approximated as the SASA of residue i in the folded structure after removing all other atoms except the main chain atoms of residues i-1 and i+1. The SASA of the folded state was measured for each atom in the context of all amyloid fibril atoms. Fibril coordinates were extended by symmetry by three to five chains on either side of the reported molecule, to ensure the energetic calculations were representative of the majority of molecules in a fibril, rather than a fibril end. To account for energetic stabilization of main chain hydrogen bonds, the ASP for backbone N/O elements was reassigned from −9 to 0 cal/mol/Å^2^ if they participated in a hydrogen bond. Similarly, if an asparagine or glutamine side chain participated in a polar ladder (two hydrogen bonds per amide) and was shielded from solvent (SASAfolded < 5 Å^2^), the ASPs of the side chain N and O elements were reassigned from −9 to 0. Lastly, the ASP of ionizable atoms (e.g. Asp, Glu, Lys, His, Arg, N-terminal amine, or C-terminal carboxylate) were assigned the charged value (−37 or −38) unless the atoms participated in a buried ion pair, defined as a pair of complementary ionizable atoms within 4.5 Å distance of each other, each with SASAfolded < 50 Å^2^). In that case, the ASP of the ion pair was reassigned to - 9*(distance - 2.8 Å)/2.8 Å)^2^. Side chain conformational entropy terms adapted from Koehl & Delarue, 1994 were added to the energy values obtained above^18^. The entropy terms were scaled by the percentage of side chain surface area buried in the assembly.

#### Construction of rIAPP homology model from hIAPP fibril structure

We modeled the six rodent substitutions on our hIAPP fibril structure using COOT^11^. We chose rotamers that minimize steric overlap for each substitution. We used Rosetta energy minimization^19^ on this model, first allowing only side chain movements (shake and wiggle) and then allowing both side chain and backbone movements. As a control, we also performed the same operations on the hIAPP fibril structure.

#### Structural alignment of hIAPP fibril structure and previously reported <u>Aβ structures</u>

PDB IDs of Aβ structures used for structural alignment are: 6OIZ^20^, 2M4J^21^, 2MVX^22^, 5KK3^23^, 5OQV^24^, 2NAO^25^, 2MXU^26^, 2BEG^27^, 2LMN^28^, 2MPZ^29^, and 6SHS^30^. Structural alignment was done via Pymol^31^. We first aligned each Aβ structure with our hIAPP fibril structure using all residues in Aβ and hIAPP, and then performed an alternative alignment using only residues 24-34 in each Aβ structure and residues 19-29 in hIAPP fibril structure. The r.m.s.d. of each alignment is summarized in Supplementary Table 3.

### Inhibitor design and model generation

Inhibitors were designed based on the same capping strategy reported previously^32^. The native structure was used as a starting point, and we selected two nine-residue segments, ^21^NNFGAILSS^29^ (N9S) and ^25^AILSSTNVG^33^ (A9G), to target inhibitors because they both have enriched β-strand conformations (5 out of 9 residues and 4 out of 9 residues, respectively) and they both contain two residues involved in the three-residue hydrophobic core that stabilizes the fibril core (Phe23 and Ile26 for N9S, and Ile26 and Val32 for A9G). To inhibit fibril growth, we selected the main chain amide-nitrogen involved in the β-sheet hydrogen bond and added a methyl group to prevent the binding of new layers by the disruption of the backbone hydrogen bonding that mediates the stacking of layers in the fibril. In addition to N9S and A9G that only target one protofilament, we selected the ^21^NNFG^24^ (N4G) segment from both protofilaments that, if considered as a continuous segment, contains more enriched β-strand content (6 out of 8 residues) than the inhibitor targets belonging to a single protofilament and also because two residues of N4G are involved in the hydrophobic core (Phe23 from both protofilament). This design is based on the observation that Gly24 from both protofilaments are very close to each other, and the direction of two N4Gs are approximately in line so they should form a long β-strand. Since two N4G segments from both protofilaments are facing each other, with the N-termini facing away from and the C-termini facing towards the center, to avoid making a bi-directional peptide, we flipped the orientation of one N4G (from Asn-Asn-Phe-Gly to Gly-Phe-Asn-Asn) and changed these flipped residues to D-amino acids. Thus the flipped and chirality-inverted N4G segment of the inhibitor should bind to hIAPP fibrils in a manner similar to the original N4G, and therefore the resulting mixed peptide can bind to both protofilaments at the same time. Since glycine does not have L- or D-chirality, the final sequence of this design is NNFGG{d-F}{d-N}{d-N} where {d-F} stands for D-phenylalanine and {d-N} stands for D-asparagine, and we named it N4Gm since it merges two N4G segments with L- and D-amino acids. We further designed the N-methylated version of N4Gm similar to N9S and A9G. The sequences of the inhibitor candidates are summarized in Supplementary Table 4. The strategy of segment selection is shown in Supplementary Fig. 6. We predict the selected segments (without N-methylation) should bind to both ends of the fibrils, whereas the designed methyl group should only allow the inhibitors to bind one end of the fibrils. This is the disadvantage of the N-methylation strategy and perhaps explains the low efficiency of designed N-methylated inhibitors.

To demonstrate the structural compatibility of our inhibitor designs with our hIAPP cryo-EM structure, we built models of the inhibitors capping the hIAPP fibrils. We manually modified one layer of the cryo-EM structure by trimming off the residues not included in the inhibitor design and adding the methyl group on the designated amide-nitrogen atom with COOT^11^. Bond length and angles were inspected to make ensure ideal values. For N4Gm inhibitors that have both L- and D-amino acids, we first took the coordinates of ^21^NNFG^24^ residues from the same layer of both protofilaments, and mutated residues from one protofilament to the opposite hand to generate the D-amino acid residues. We renumbered the D-amino acid residues to be consecutive after the L-amino acid residues in COOT^11^, and we used CNS^33^ to minimize the energy. The methyl group of N4Gm inhibitors was also added as previously described.

### ThT assays

Synthetic hIAPP wild type or S20G peptide was diluted to 10 μM in PBS supplemented with 30 μM ThT and filtered using 0.1 μm Ultrafree-MC-VV centrifugal filters (Millipore). Filtered solution was mixed with 100:1 (v/v) 100% DMSO, 30 mM inhibitor dissolved in DMSO, or 3 mM inhibitor dissolved in DMSO, resulting in 10 μM hIAPP alone, 10 μM hIAPP with 30 μM inhibitor, or 10 μM hIAPP with 300 μM inhibitor. PBS with the same concentration of ThT and DMSO was used as buffer control. The sample solution was pipetted into a polybase black 384-well plate with optical bottom (Thermo Scientific) and incubated at 37 °C without shaking. ThT fluorescence was measured with excitation and emission wavelengths of 440 and 480 nm, respectively, using FLUOstar Omega plate reader (BMG LABTECH). The aggregation curves were averaged from three to four independent measured replicates and error bars show s.d. of replicate measurements. To normalize the different ranges of fluorescence readings observed from each experiments (probably due to the different fluorescence gain settings of the plate reader), we normalized the readings to make the minimum mean value in each panel 0% and the maximum mean value in each panel 100%.

## Supplemental Figures

**Supplementary Figure 1.**
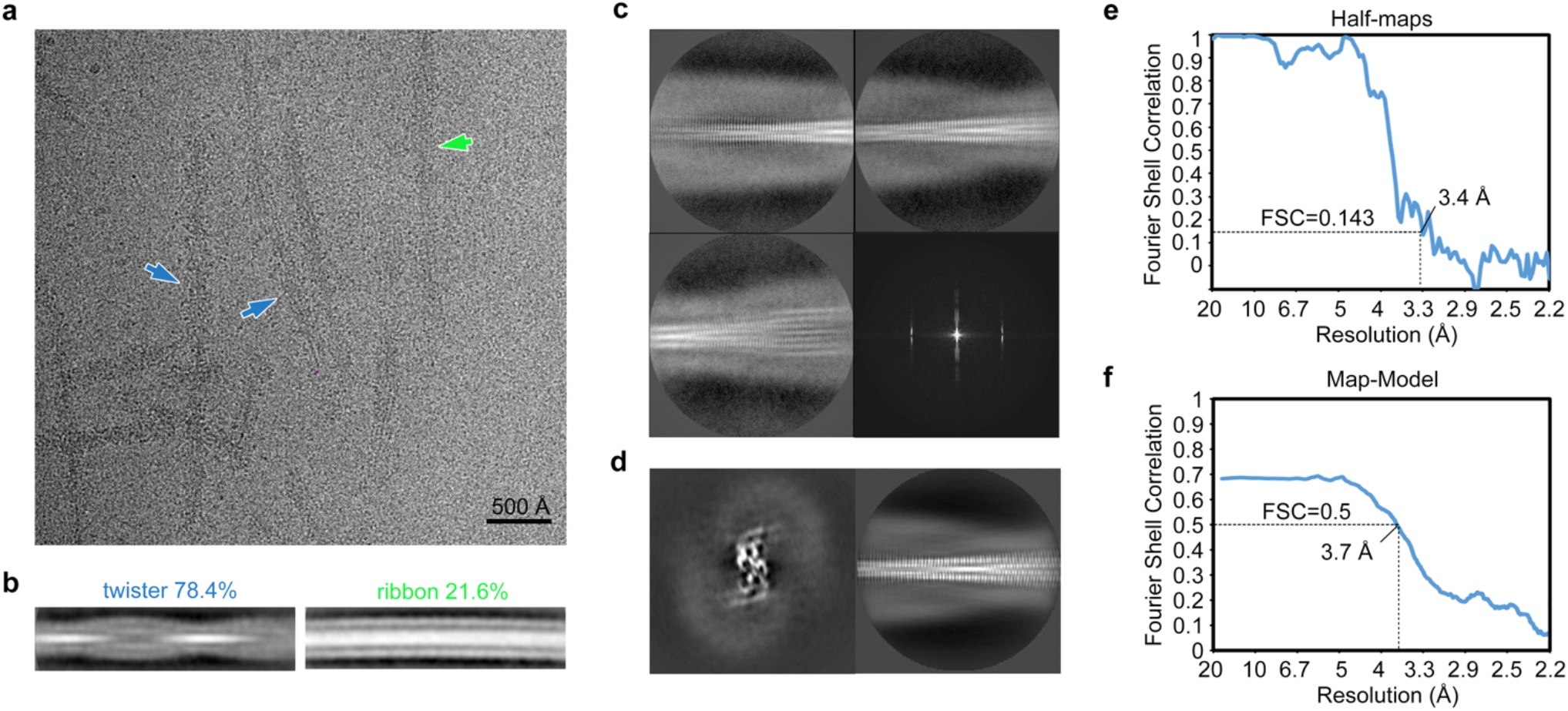
Cryo-EM data processing. **a,** Representative Krios micrograph of hIAPP fibrils. Blue and green arrows indicate two morphologies (twister and ribbon, respectively) identified by 2D classification. Notice they are not distinguishable by eye. **b,** Representative 2D classes and relative population of twister and ribbon morphologies. **c,** Representative 2D classes of twister with smaller box size particles showing the 4.8 Å β-sheet spacing, and the computed diffraction pattern from a representative 2D class. **d,** Central slice (left) and 2D projection (right) from the final reconstruction. Notice the 2D projection of final reconstruction is consistent with 2D classification. **e-f,** FSC curves between two half-maps (**e**) and the cryo-EM reconstruction and refined atomic model (**f**).

**Supplementary Figure 2.**
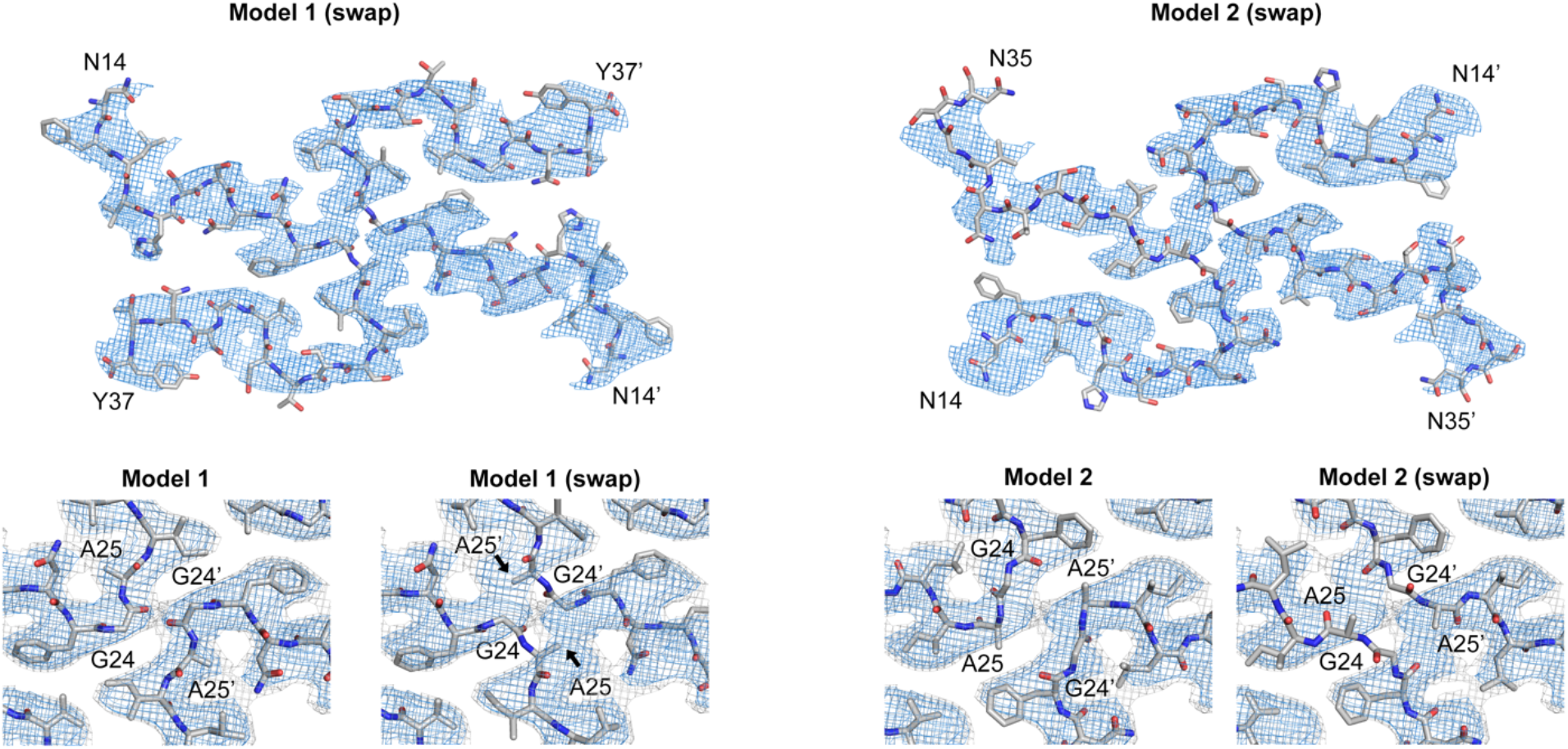
Potential domain swapping of hIAPP models. Domain swapped versions of both Model 1 and Model 2 were built to test the possibility of domain swapping. In the swapped models, the residues between the N-terminus and Gly24 from one protofilament were connected to the residues between Ala25 and the C-terminus from the other protofilament of the un-swapped model. The density map with σ=3.0 is shown in blue mesh and that with σ=2.0 is shown in grey mesh. Notice that Gly24 in both swapped Model 1 and Model 2 is clearly out of the density, demonstrating that the domain swapping is not supported by our cryo-EM map.

**Supplementary Figure 3.**
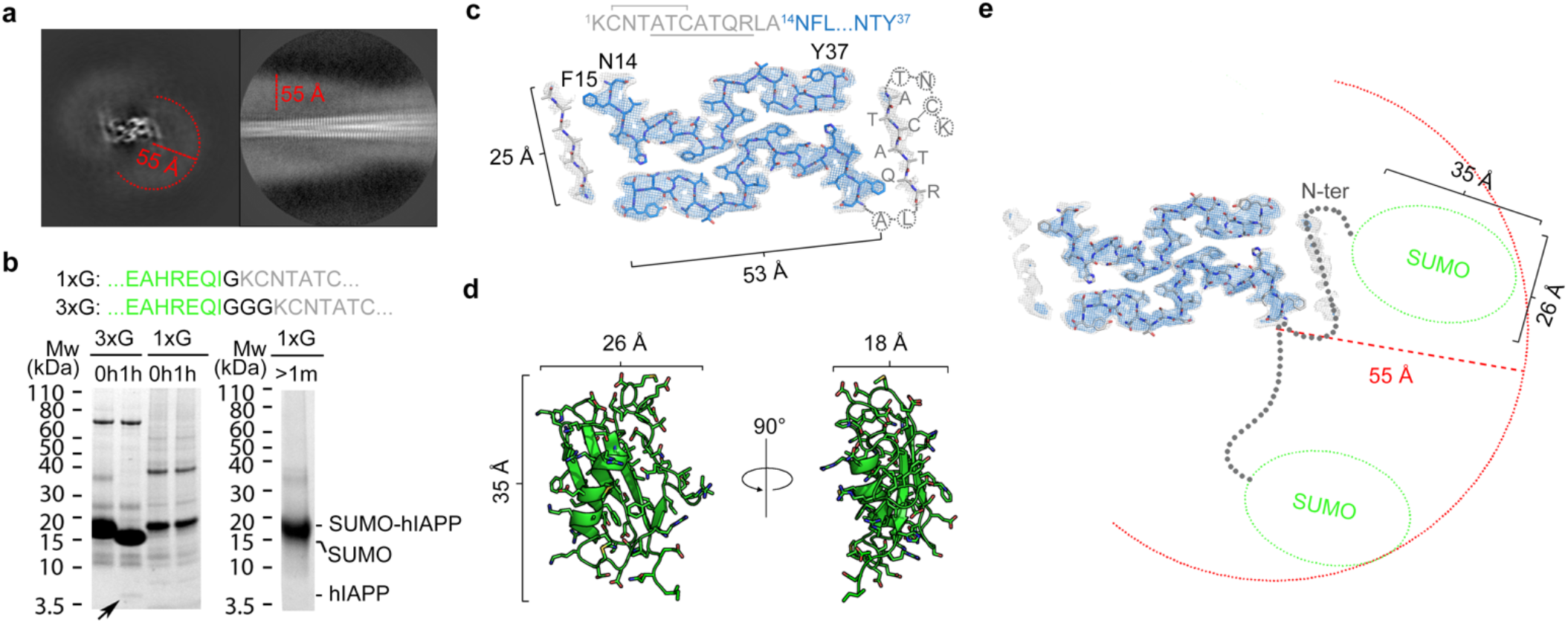
The fuzzy coat in hIAPP fibril structure may represent the flexible N-terminal of hIAPP and the SUMO-tag. **a,** The final reconstruction (left) and 2D classification (right) show a fuzzy coat of ~55 Å surrounding the fibril core. **b,** Protease cleavage assays indicate the construct we used for fibril structure determination (SUMO-IAPP with 1xG, means one glycine between SUMO-tag and hIAPP) has an un-removable SUMO-tag, whereas the SUMO-tag is removable when we extend the linker to three glycine. **c,** Plausible N-terminal conformation suggested by the extra densities near Asn14. The density map with σ=3.0 is shown in blue mesh and that with σ=2.0 is shown in grey mesh. The intra-molecular disulfide bond is labelled between Cys2 and Cys7, and the residues occupying the extra densities in our hypothetical model are underlined. **d,** Crystal structure of SUMO protein (PDB ID 1L2N). **e,** Hypothetical model of N-terminus of hIAPP and SUMO-tag match the dimensions of the fuzzy coat observed in the hIAPP fibril reconstruction. Notice that in most cases the SUMO-tag is far away from the fibril core therefore should not influence the fibril structure.

**Supplementary Figure 4.**
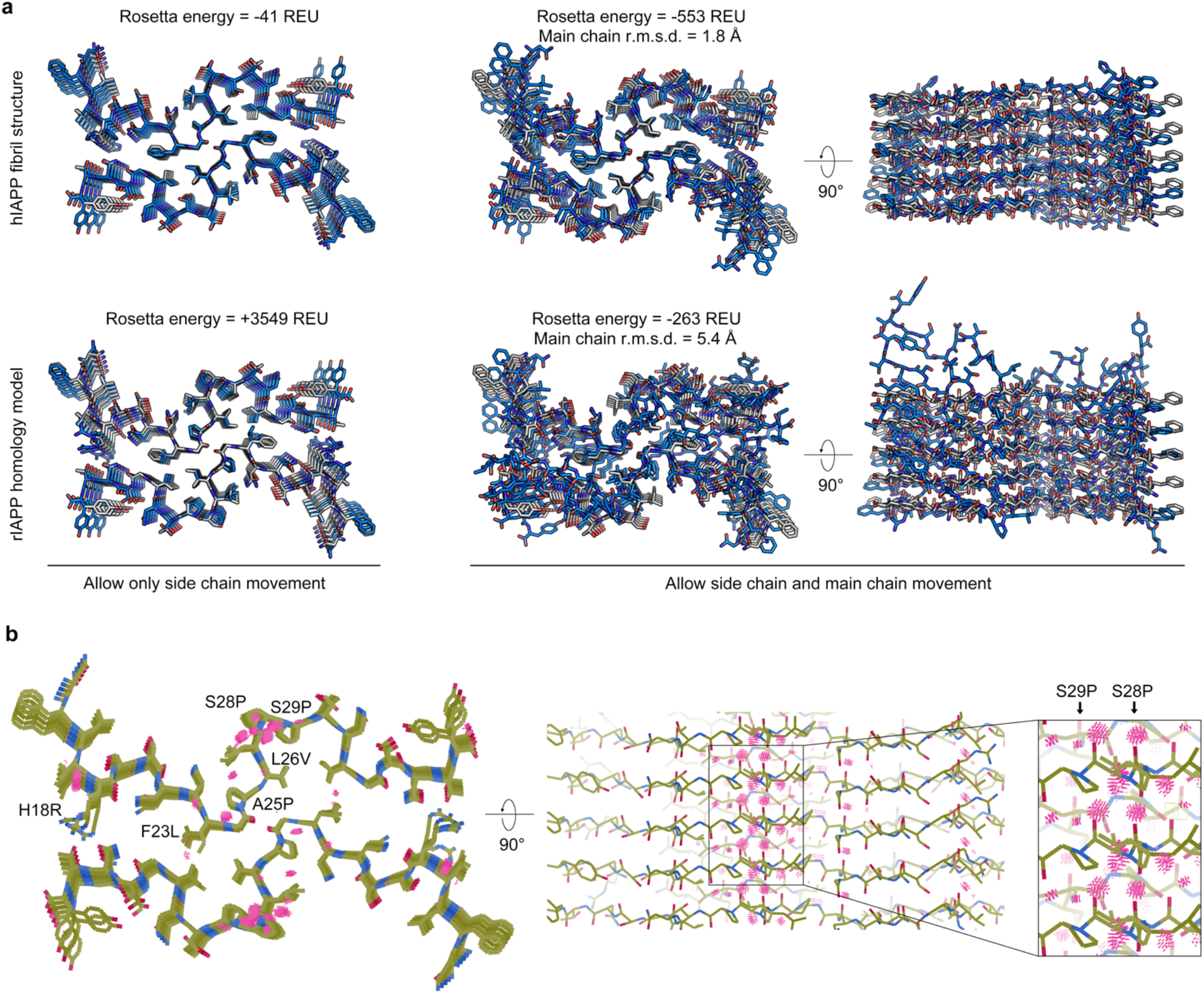
Rosetta energy minimization of hIAPP fibril structure and rIAPP homology model. **a,** Structure superimposition between (grey) hIAPP fibril structure determined here and (blue) hIAPP fibril structure (upper panels) or rIAPP homology model (lower panels) optimized by Rosetta energy minimization. Calculation was done either allowing only side chain movements (left panels) or allowing both side chain and main chain movements (middle and right panels). Notice that during Rosetta energy minimization, we did not apply non-crystallographic symmetry so that the 5 layers in each model were not forced to be identical. **b,** Steric clashes of the rIAPP homology model after side chain Rosetta energy minimization were probed with COOT and displayed as red dots. Notice that most of the steric clashes are found near S28P and S29P.

**Supplementary Figure 5.**
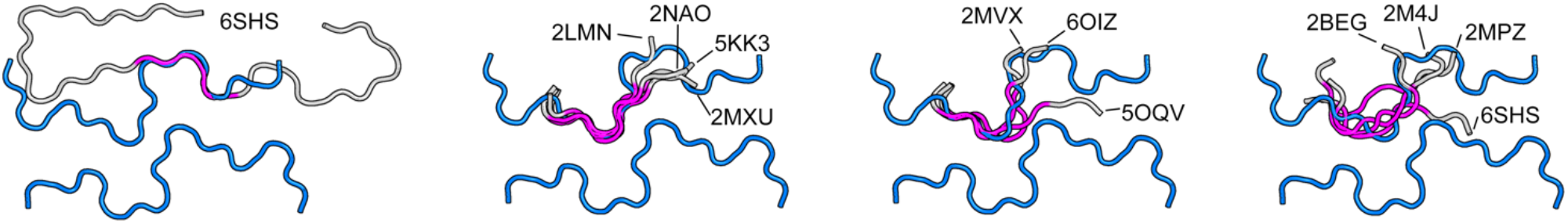
Structural superimposition of Aβ fibril structures and hIAPP fibril structure. Ten previously reported Aβ fibril structures were superimposed with the hIAPP fibril structure by either directly comparing full-length Aβ fibril structures with the full-length hIAPP structure, or by only comparing residues 24-34 of Aβ fibril structures with residues 19-29 of the hIAPP structure. For the full-length comparison, one Aβ fibril structure (PDB ID 6SHS) shows reasonable alignment with low r.m.s.d., and the structural superimposition is shown on the far left panel, with the Aβ fibril structure shown in grey, the hIAPP structure shown in blue, and the segment that fits best (residues 20-25 of Aβ fibril structure) shown in magenta. For the partial comparison, four Aβ fibril structures show a good fit (middle left), three Aβ fibril structures show a moderate fit (middle right) and four Aβ fibril structures do not fit (far right). In these superimpositions, residues 24-34 of the Aβ fibril structures were colored grey and the highest fitting region (residues 26-31) is colored magenta. Detailed alignment parameters are listed in Supplementary Table 3.

**Supplementary Figure 6.**
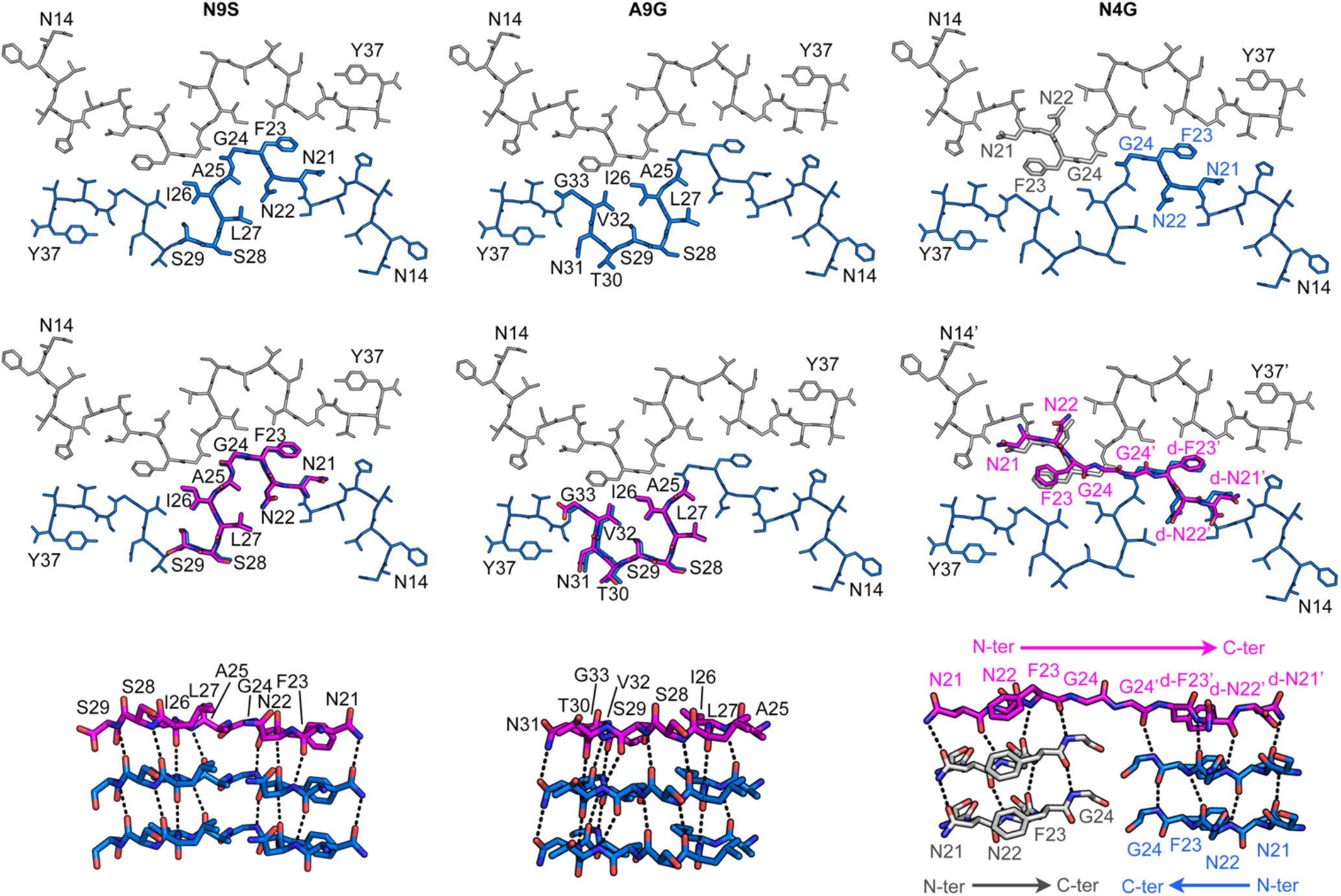
Segments selected for hIAPP fibril inhibitor design. Three segments of hIAPP, ^21^NNFGAILSS^29^ (N9S, left panels), ^25^AILSSTNVG^33^ (A9G, middle panels) and ^21^NNFG^24^ (N4G, right panels), were selected for design of inhibitors of hIAPP fibrils. For each selected segment, the hIAPP structure with the segment highlighted is shown on the top, with the hIAPP structure shown as lines and the segment shown as sticks. Proposed models of the corresponding inhibitor peptides (before adding N-methylation) binding to the hIAPP structures are shown as top views (middle panels) and side views (bottom panels). Notice there are multiple hydrogen bonds between the designed inhibitors and hIAPP fibrils, providing binding affinities for these inhibitors. For the N4G merged inhibitor, the model indicates the orientation-flipped and chirality-reversed N4G has high structural similarity to the original N4G and recaptures all original inter-layer interactions. Hydrogen bonds with distances between 2.3-3.2 Å are shown as black dashed lines.

**Supplementary Figure 7.**
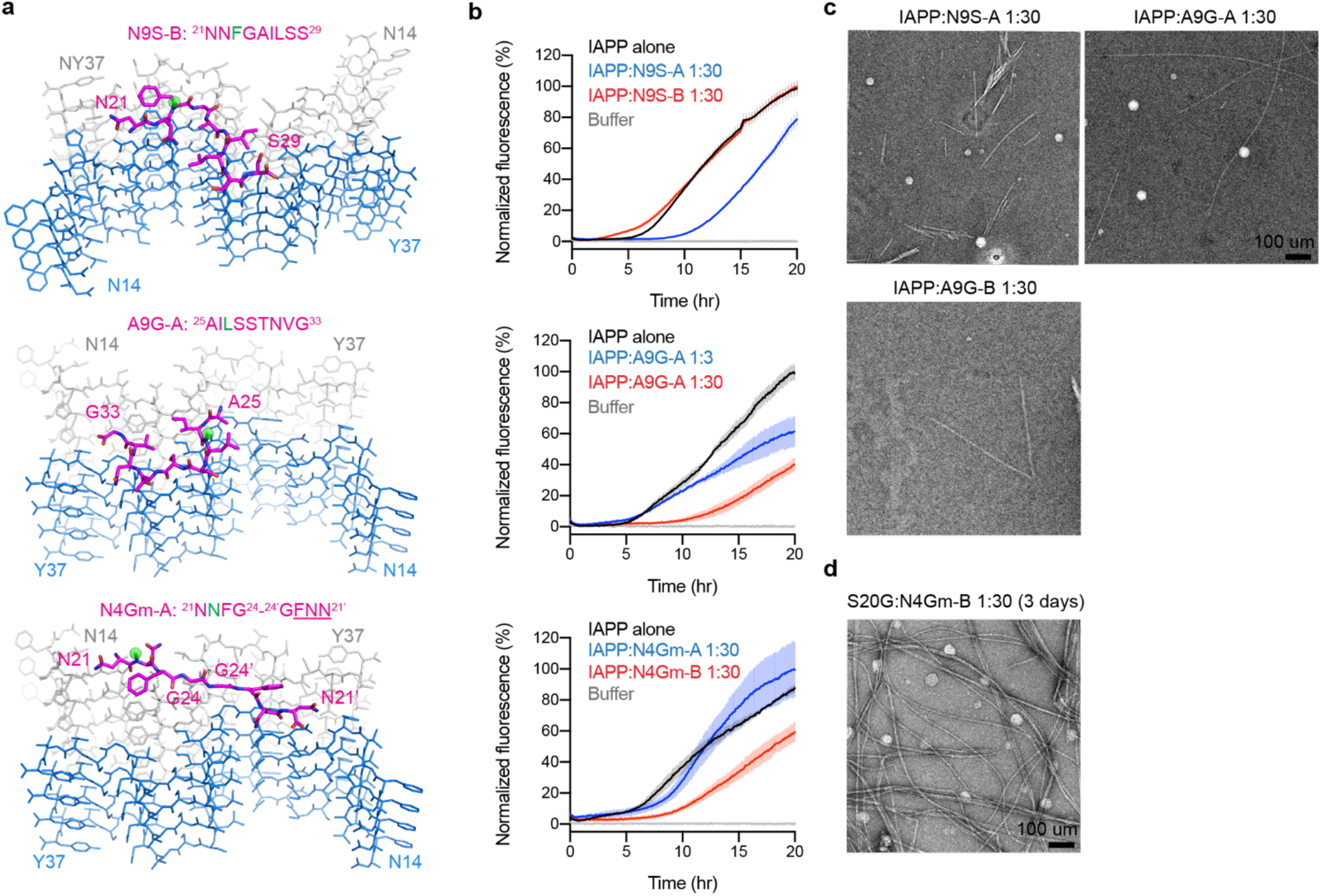
Additional inhibitors designed for hIAPP fibrils. **a,** Proposed model of designed inhibitors (magenta) bound to hIAPP fibrils (blue and grey for each protofilament). The methyl group of N-methylated inhibitors is shown as a green sphere. The last three residues of N4Gm-A are d-amino acids and are underlined. **b,** ThT assays measuring inhibitor efficacy shown on the left. A9G-A delays hIAPP fibril formation but not N9S-B and N4Gm-A. For the two inhibitors that were not effective, two effective inhibitors (N9S-A and N4Gm-B, respectively) are tested in the same experiment as controls. Data are shown as mean ± s.d., n=3 independent experiments. **c,** Negative stain EM images of hIAPP with N9S-A, A9G-A or A9G-B after 20 hours of incubation. Notice that hIAPP fibril formation is not fully eliminated by these inhibitors. **d,** Negative stain EM shows hIAPP S20G fibrils present after 3 days of incubation with N4Gm-B, suggesting that fibril formation of hIAPP S20G is not fully eliminated when longer incubation times are examined (compared to 20 hours shown in Fig. 3e).

**Supplementary Figure 8.**
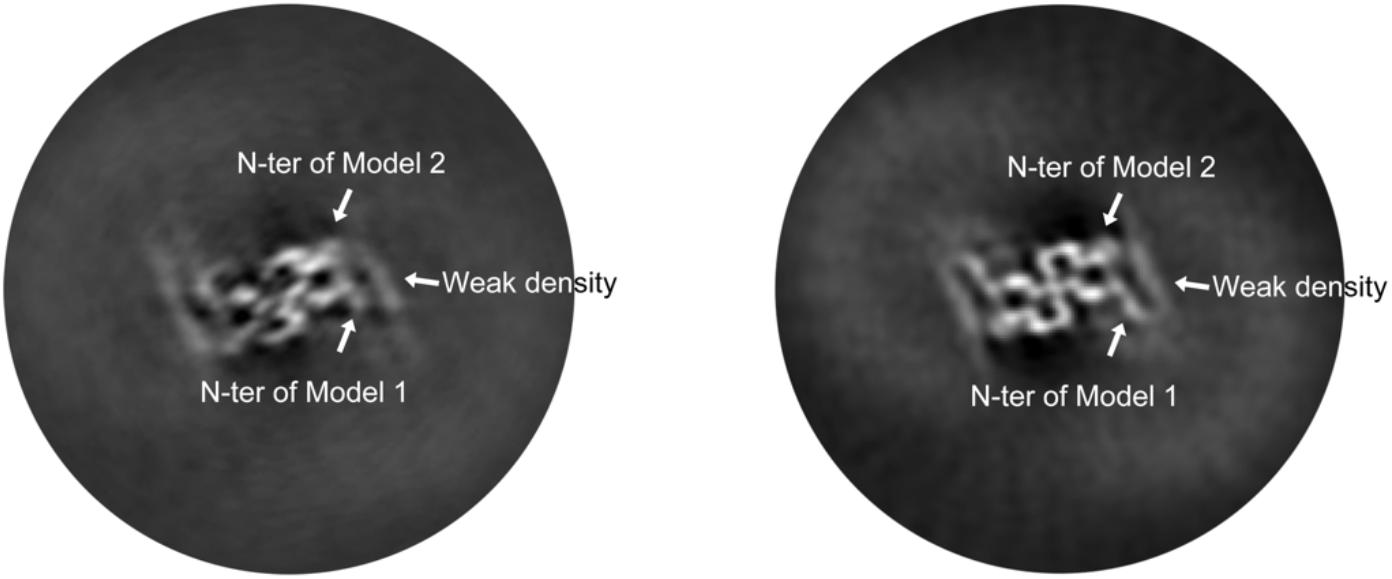
Connection of N-terminal density. Slices of 3D maps of the final reconstruction (left) and an earlier reconstruction with lower resolution (right). The positions that represent N-terminus of Model 1 and Model 2 are indicated by arrows. Note the weak density that represents the flexible N-terminus of hIAPP seems to connect to the position that represents the N-terminus of Model 2 in the final reconstruction (left); whereas in the lower resolution reconstruction (right), the weak density seems to connect to the position of N-terminus of Model 1.

## Supplemental Tables

**Supplementary Table 1.**
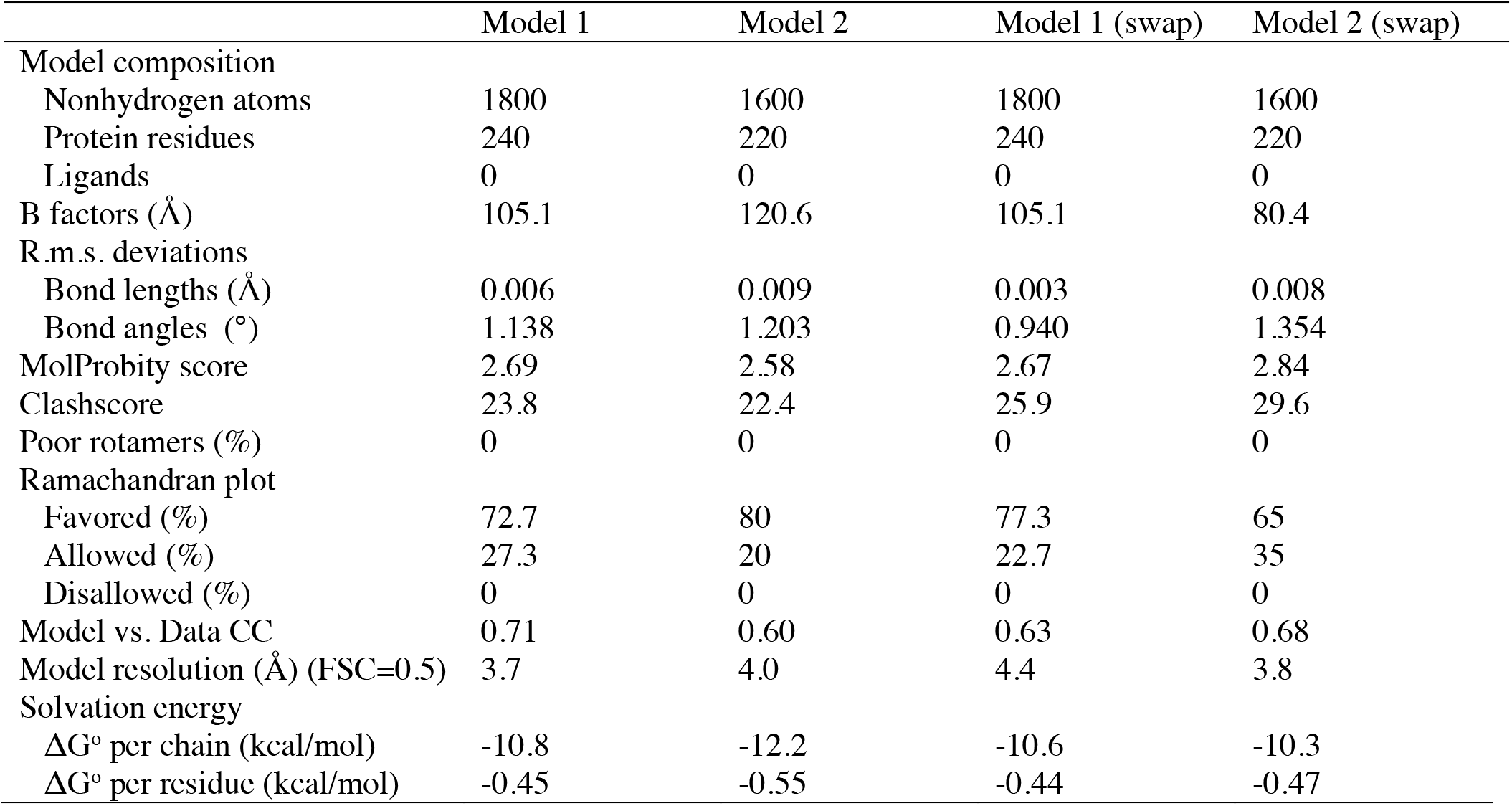
Comparison of possible models of hIAPP fibrils

|  | Model 1 | Model 2 | Model 1 (swap) | Model 2 (swap) |
| --- | --- | --- | --- | --- |
| Model composition |  |  |  |  |
| Nonhydrogen atoms | 1800 | 1600 | 1800 | 1600 |
| Protein residues | 240 | 220 | 240 | 220 |
| Ligands | 0 | 0 | 0 | 0 |
| B factors (Å) | 105.1 | 120.6 | 105.1 | 80.4 |
| R.m.s. deviations |  |  |  |  |
| Bond lengths (Å) | 0.006 | 0.009 | 0.003 | 0.008 |
| Bond angles (°) | 1.138 | 1.203 | 0.940 | 1.354 |
| MolProbity score | 2.69 | 2.58 | 2.67 | 2.84 |
| Clashscore | 23.8 | 22.4 | 25.9 | 29.6 |
| Poor rotamers (%) | 0 | 0 | 0 | 0 |
| Ramachandran plot |  |  |  |  |
| Favored (%) | 72.7 | 80 | 77.3 | 65 |
| Allowed (%) | 27.3 | 20 | 22.7 | 35 |
| Disallowed (%) | 0 | 0 | 0 | 0 |
| Model vs. Data CC | 0.71 | 0.60 | 0.63 | 0.68 |
| Model resolution (Å) (FSC=0.5) | 3.7 | 4.0 | 4.4 | 3.8 |
| Solvation energy |  |  |  |  |
| ΔG° per chain (kcal/mol) | -10.8 | -12.2 | -10.6 | -10.3 |
| ΔG° per residue (kcal/mol) | -0.45 | -0.55 | -0.44 | -0.47 |

**Supplementary Table 2.**
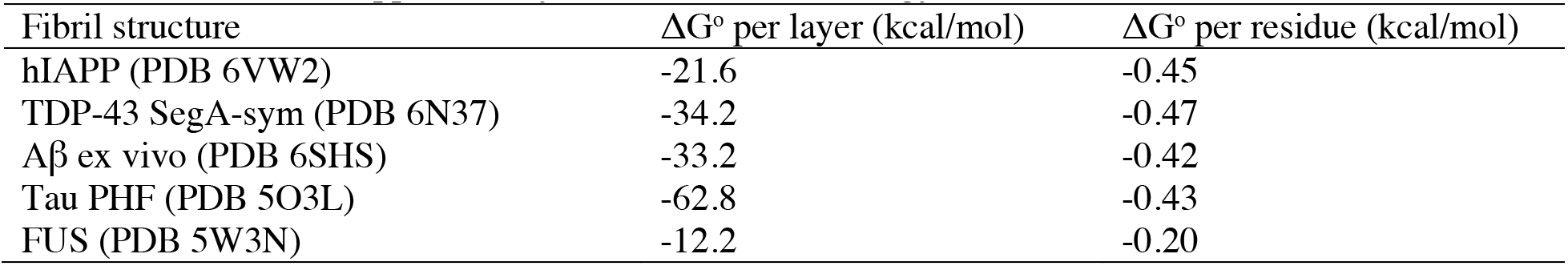
Solvation energy calculation

**Supplementary Table 3.** Structure alignments between Aβ and hIAPP

| Aβ structures used for alignment |  |  |  | Full length alignment |  | IAPP 24-34 vs. Aβ 19-29 |  |
| --- | --- | --- | --- | --- | --- | --- | --- |
| PBD ID | Residues | Mutation | Methods | r.m.s.d. (Å) | # of atoms | r.m.s.d. (Å) | # of atoms |
| 6OIZ | 20-34 | isoAsp23 | MicroED | 4.0 | 42 | 2.1 | 40 |
| 2M4J | 1-40 | Wild type | NMR | 3.3 | 55 | 3.5 | 42 |
| 2MVX | 1-40 | Osaka (E22Δ) | NMR | 6.4 | 81 | 1.5 | 36 |
| 5KK3 | 1-42 | Wild type | NMR | 4.7 | 62 | 1.5 | 38 |
| 5OQV | 1-42 | Wild type | CryoEM | 4.8 | 62 | 1.9 | 36 |
| 2NAO | 1-42 | Wild type | NMR | 4.8 | 61 | 1.6 | 40 |
| 2MXU | 1-42 | Wild type | NMR | 4.9 | 62 | 1.6 | 39 |
| 2BEG | 1-42 | Wild type | NMR | 3.7 | 55 | 2.0 | 40 |
| 2LMN | 1-40 | Wild type | NMR | 3.5 | 59 | 1.8 | 42 |
| 2MPZ | 1-40 | Lowa (D23N) | NMR | 3.7 | 56 | 2.8 | 42 |
| 6SHS | 1-40 | Wild type (ex vivo) | CryoEM | 1.8 | 46 | 2.3 | 35 |

**Supplementary Table 4.**
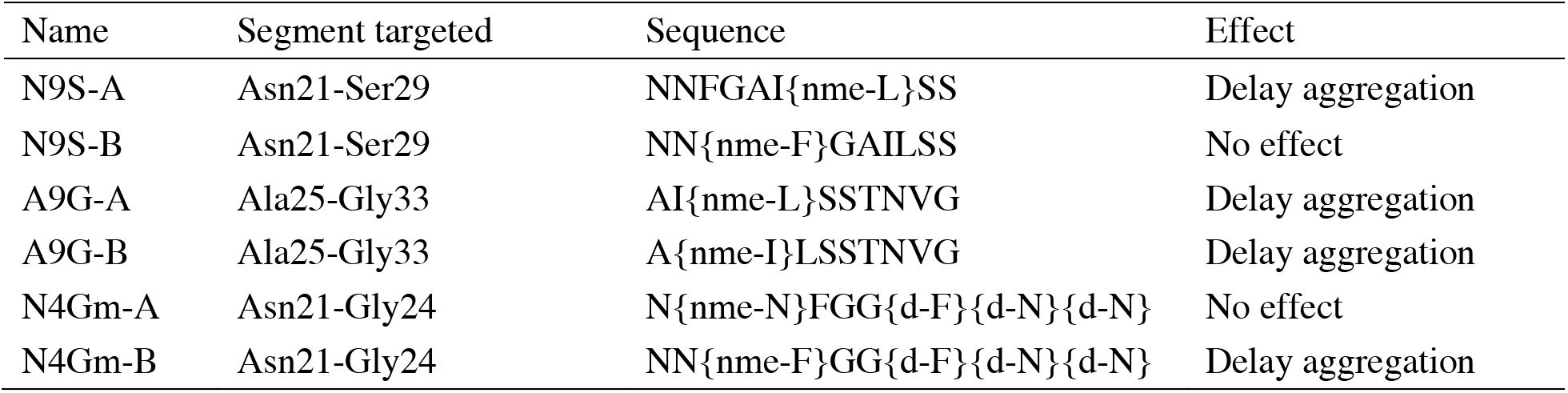
Inhibitors designed based on hIAPP cryo-EM structure

